# Enhanced lipogenesis through Pparγ helps cavefish adapt to food scarcity

**DOI:** 10.1101/2021.04.27.441667

**Authors:** Shaolei Xiong, Wei Wang, Alexander Kenzior, Luke Olsen, Jaya Krishnan, Jenna Persons, Kyle Medley, Robert Peuß, Yongfu Wang, Shiyuan Chen, Ning Zhang, Nancy Thomas, John M. Miles, Alejandro Sánchez Alvarado, Nicolas Rohner

## Abstract

Nutrient availability varies seasonally and spatially in the wild. The resulting nutrient limitation or restricted access to nutrients pose a major challenge for every organism. While many animals, such as hibernating animals, evolved strategies to overcome periods of nutrient scarcity, the cellular mechanisms of these strategies are poorly understood. Cave environments represent an extreme example of nutrient deprived environments since the lack of sunlight and therefore primary energy production drastically diminishes the nutrient availability. Here, we used *Astyanax mexicanus*, which includes river-dwelling surface fish and cave adapted cavefish populations to study the genetic adaptation to nutrient limitations. We show that cavefish populations store large amounts of fat in different body regions when fed ad libitum in the lab. We found higher expression of lipogenesis genes in cavefish livers when fed the same amount of food as surface fish, suggesting an improved ability of cavefish to use lipogenesis to convert available energy into triglycerides for storage into adipose tissue. Moreover, the lipid metabolism regulator, Peroxisome proliferator-activated receptor γ (Pparγ), is upregulated at both transcript and protein levels in cavefish livers. Chromatin Immunoprecipitation sequencing (ChIP seq) showed that Pparγ binds cavefish promoter regions of genes to a higher extent than surface fish. Finally, we identified two possible regulatory mechanisms of Pparγ in cavefish: higher amounts of ligands of the nuclear receptor, and nonsense mutations in *per2*, a known repressor of Pparγ. Taken together, our study reveals that upregulated Pparγ promotes higher levels of lipogenesis in the liver and contributes to higher body fat accumulation in cavefish populations, an important adaptation to nutrient limited environments.

## Introduction

Nutrient availability can vary greatly throughout the year. To adapt to periods of dearth, most animals will store excess energy in the form of fat when food is available and utilize these fat storages when food is scarce. Such fat gains can be impressive. Brown bears can gain up to 180 kg of weight, most of it as fat, in the few summer months before hibernation (Kingsley et al., 1983), and migrating birds can build up fat stores that make up to 50% of their bodyweight before migrating (Blem, 1976). Other extreme examples are found in cave animals. Due to the lack of sunlight and therefore primary production, cave habitats rely on nutrients that originate outside of the caves and are transported only occasionally into the cave through floods or bat droppings (Mitchell et al., 1977). One well studied example is the teleost species, *Astyanax mexicanus*. Previous studies have shown that cavefish populations of this species can gain substantially higher amounts of total body fat and visceral fat compared to the surface forms of the same species (Aspiras et al., 2015; Hüppop, 2001; Xiong et al., 2018). Some cavefish populations (*e.g*., Tinaja or Molino) achieve fat gain through hyperphagia caused by nonsynonymous mutations in the melanocortin 4 receptor (*mc4r*) (Aspiras et al., 2015). Notably, the same mutations cause hyperphagia in humans (Aspiras et al., 2015). However, not all cavefish populations carry these mutations or display increased appetite. For example, cavefish from the Pachón population carry the wildtype allele of *mc4r* and display comparable appetites to surface fish (Alie et al., 2018; Aspiras et al., 2015). Several strategies have been proposed to explain how Pachón gain high body fat. Pachón develop visceral adipocytes earlier than surface fish (Xiong et al., 2018), their gastrointestinal tract has higher digestion/absorption efficiency than surface fish (Riddle et al., 2018), and their skeletal muscle are insulin resistant, a proposed adaptation to the nutrient-limited environment (Riddle et al., 2018b). However, it is not known whether there are specific cellular changes to Pachón’s lipid metabolism.

A central metabolic pathway that controls the synthesis of lipids through fatty acid biosynthesis and triglyceride production in liver and adipose tissue is lipogenesis (Kersten, 2001; Numa and Yamashita, 1974; Wang et al., 2015). In this study, we investigated whether the observed differences in body fat content between cavefish and surface fish of *A. mexicanus* are related to changes in lipogenesis. We observed that Pachón cavefish display a massive transcriptional upregulation of central genes in the lipogenesis pathway in the liver after feeding (up to 100-fold), compared to surface fish. This was accompanied by an upregulation of a central regulator of lipogenesis, the transcription factor Peroxisome proliferator-activated receptor gamma (Pparγ). Moreover, we found increased levels of Pparγ activators and mutations in a direct repressor of Pparγ in cavefish populations, supporting the notion that higher activity of lipogenesis through Pparγ transcriptional and functional upregulation underlies the increased adipogenesis observed in cavefish.

## RESULTS

### Cavefish display increased body fat levels throughout the body

Previous studies have shown that compared to surface fish, cavefish populations display higher total triglycerides and visceral fat when fed ad libitum (Aspiras et al., 2015; Hüppop, 2001; Xiong et al., 2018). To confirm these findings and develop a method to more easily quantify total body fat in fish, we used EchoMRI to measure body fat percentage. Consistent with previous total triglycerides measurements (Aspiras et al., 2015), fish from both the Pachón and Tinaja populations showed higher body fat than surface fish (mean surface=15.2%; mean Pachón=39.6%; mean Tinaja=34.2%; Fig. 1A). To better visualize fat distribution throughout the body, we dissected adult fish into various sections and used hematoxylin and eosin (H&E) staining of head and trunk sections. We observed that Tinaja and Pachón cavefish store fat in the entire eye socket which is absent in surface fish (Fig. 1B). Tinaja and Pachón cavefish have markedly more adipocytes in the ventral part and lateral sides of the trunk compared to surface fish (Fig. 1C). In the dorsal part of the trunk, we observed only slightly more adipocytes in cavefish compared to surface fish (Fig. 1C). In total, the relative adipose area in the transverse trunk section of cavefish was significantly higher than that of surface fish (Fig. 1D). Additionally, we used the Folch method, which takes advantage of the biphasic solvent system consisting of chloroform/methanol/water (Folch et al., 1957) to extract and quantify total lipid content from head, dorsal and ventral parts of the trunk (Fig. 1E). We found Tinaja and Pachón cavefish have higher lipid content in the head, dorsal trunk and ventral trunk sections compared to surface fish (Fig. 1F-H). Notably, we found no difference in hepatic triglyceride and total liver lipid between adult surface fish and cavefish populations (Supplementary Fig. 1), indicating that one-year old cavefish do not over accumulate lipid in the liver, at least on our lab feeding regime (see methods for detail).

**Fig 1.**
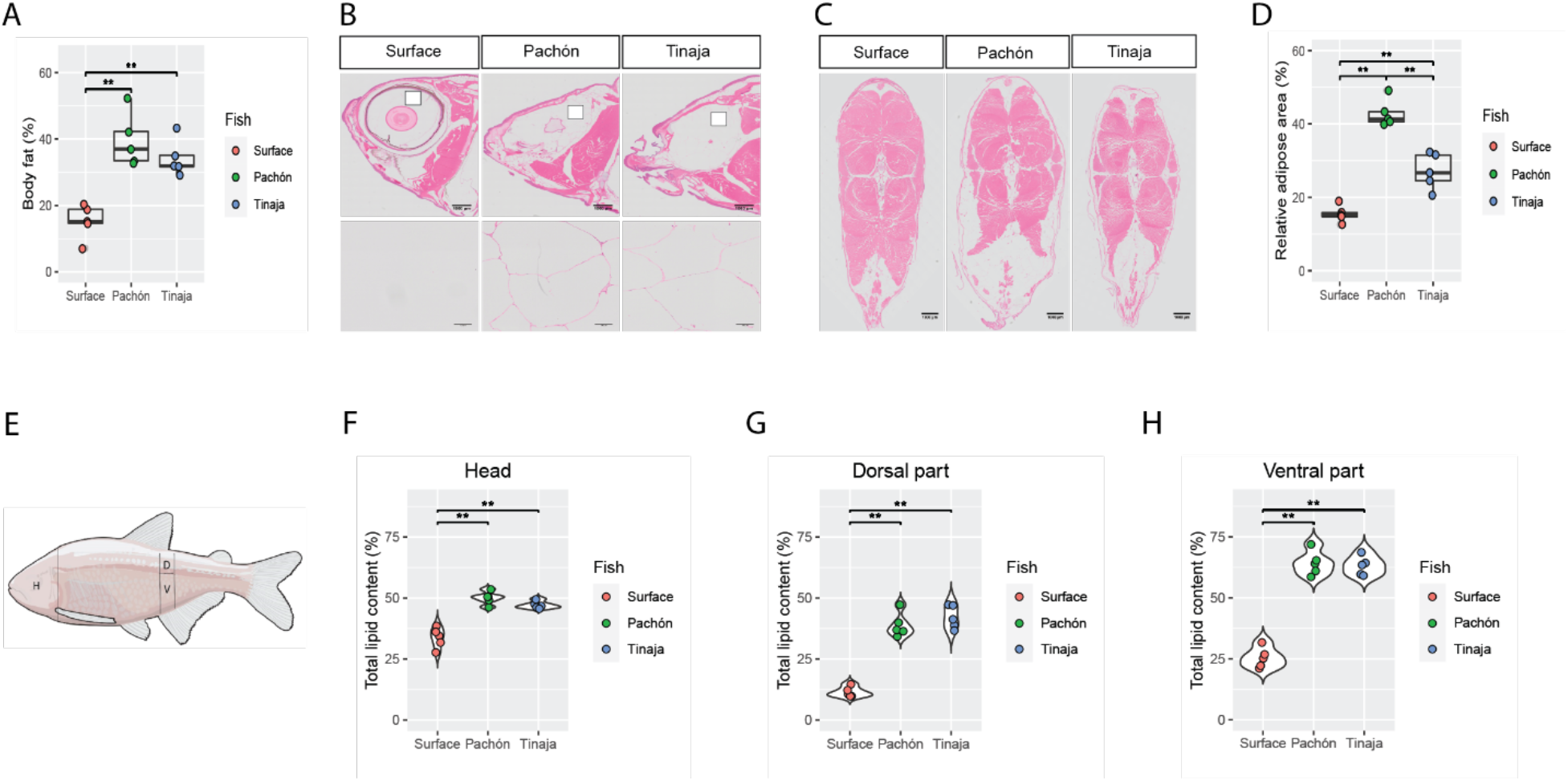
Cavefish display more body fat in various areas of the body compared to surface fish. A) Total body fat comparison (fat mass/total body weight) between surface fish, and Pachón and Tinaja cavefish using EchoMRI (n=5 per population). B) H&E staining of fish head sections of the three fish populations (Surface, Pachón, Tinaja). The sagittal sections were performed across the eye area of the head, the upper panel showing the entire section and the lower panel showing the region indicated with a white box in the upper panel, revealing that in cavefish the eye socket is filled with adipocytes (n=5 per population, scale bar=1 mm in the upper panel, scale bar=100 μm in the bottom panel). C) Transverse H&E staining of fish trunk sections close to the anal fin of the three fish populations (n =5 per population, scale bar =1 mm). D) Quantification of fat area to the whole transverse trunk section area in surface fish, Pachón, and Tinaja cavefish using “Convert to Mask” in ImageJ (n=5 per population). E) Cartoon highlighting sampling areas for total lipid content quantification (H = head, D = dorsal part, V = ventral part, black lines indicate the boundaries of sampling). F-H) Total lipid content (%) in surface fish, Pachón, and Tinaja cavefish (n=5 per population) using the Folch method. Significances calculated with Wilcoxon test, **p <0.01.

### Cavefish increase lipogenesis in the liver

Given the differences in body fat content between cavefish and surface fish populations throughout different body parts, we hypothesized that cavefish display higher postprandial lipid anabolism than surface fish. This should be particularly pronounced in Pachón cavefish as they are known to have similar appetites as surface fish (Alie et al., 2018; Aspiras et al., 2015). To study the transcriptional response to feeding, we first fasted juvenile Pachón cavefish and surface fish for 4 days to allow transcription of anabolic genes to cease. We then refed the different populations the same amount of food and performed bulk RNA-seq of liver tissue, which is a primary center of lipogenesis (Fig. 2A). We identified ~16,000 genes (TPM > 1), of which ~2,300 were differentially expressed (DE) between the refed Pachón and surface fish samples (Supplementary Fig. 2). We performed GO-term enrichment analysis of the DE genes and identified numerous overrepresented metabolic pathways in the cavefish samples (Fig. 2B). Among these enriched terms in the cavefish samples, we identified lipid anabolic pathways such as fatty acid biosynthesis and triglyceride biosynthesis (Fig. 2B). To further dissect these pathways, we focused our analyses on key genes of these pathways (i.e., *aclya, acaca, fasn, scd, elovl6, gpam, dgat1b, dgat2, lpin1, acsl4a, oxsm, olah*) (Fig. 2C). Interestingly, these genes showed similar expression level at the fasted state between surface fish and Pachón cavefish, but much higher expression levels in refed Pachón cavefish compared to refed surface fish (Fig. 2C). These results indicate a likely enhanced postprandial lipogenic capacity within the Pachón cavefish. We confirmed these results by focusing on three key fatty acid biosynthesis genes (ATP citrate lyase (*acly*), acetyl-CoA carboxylase 1 (*acaca*), and fatty acid synthase (*fasn*)) using qRT-PCR. All three genes responded to feeding by a 10-100 fold increased expression in Pachón cavefish compared to surface fish (Fig. 2D), suggesting that Pachón cavefish have a greater ability to synthesize fatty acids following feeding than surface fish.

**Figure 2.**
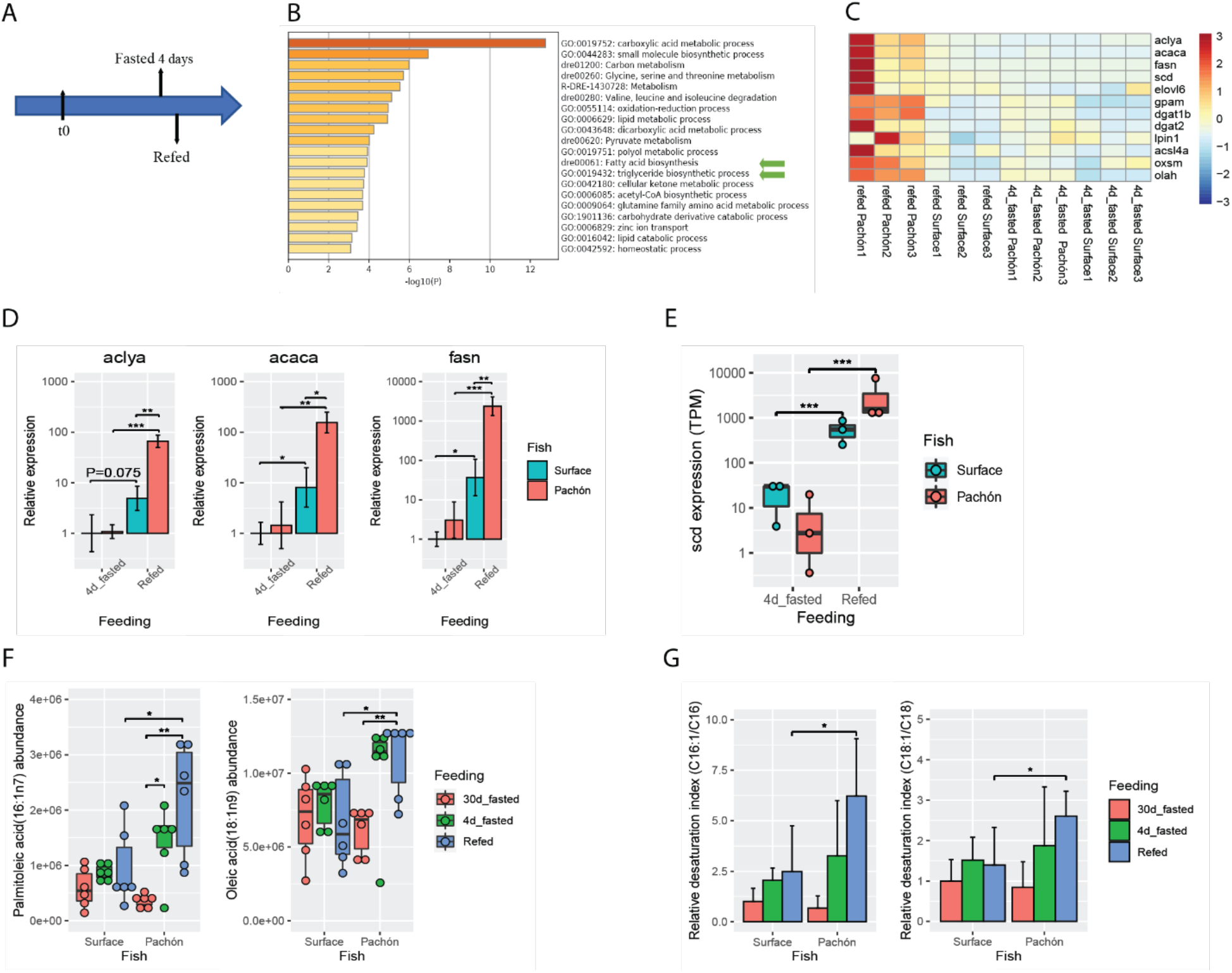
Lipogenesis genes are upregulated in the liver and fatty acid profile is altered in Pachón cavefish compared to surface fish. A) Experimental design schematic for RNA-seq analysis of Pachón and surface fish (n=3 per population and condition). B) GO-term comparison and analysis of upregulated genes in refed Pachón and surface fish livers. Green arrows indicate the key lipid anabolic pathways, fatty acid biosynthesis and triglyceride biosynthesis process. C) Heatmap of lipogenesis genes in fasted and refed surface fish and Pachón cavefish. D) Relative expression (RT-qPCR) of fatty acid biosynthesis genes in livers of fasted and refed surface fish and Pachón cavefish (n=3, t-test). E) Expression of *scd* in livers of fasted and refed surface fish and Pachón cavefish (n=3) TPM: transcript per million. F) Fatty acid profiles of two monounsaturated fatty acids (n=6 wilcox test) data from (Medley et al., 2020). G) Refed Pachón cavefish livers have a higher desaturation index (C16:1n7/C16, and C18:1n9/C18) than surface fish (n=6, wilcox test) data from (Medley et al., 2020) *p<0.05; **p<0.01; ***p<0.001.

To better understand the temporal dynamics of postprandial gene expression – specifically the duration of increased lipogenesis - we performed a time course study of lipogenic gene expression. We measured transcription levels of key fatty acid biosynthesis genes (*acly, acaca, fasn*) and triglyceride biosynthesis genes (*scd1, elovl6, gpam*, and *dgat2*) using qRT-PCR of liver tissues of surface fish and Pachón cavefish at different time points after paired feeding. We found higher expression of these lipogenesis genes in Pachón cavefish samples compared to surface fish up to 24 hours after feeding, with the highest expression at the 6-hour time point (Supplementary Fig. 3). The gene expression differences were no longer detected at the 5-day time point. This indicates that upregulation of lipogenesis can last more than 24 hours (but less than 5 days) after feeding the same amount of food in Pachón cavefish compared to surface fish.

To independently confirm whether increased lipogenesis in cavefish was occurring, we measured the conversion of saturated fatty acids to monounsaturated fatty acids, a key step in the generation of triglycerides from fatty acids (Ntambi and Miyazaki, 2004; Wang et al., 2015). The products of such conversion are monounsaturated fatty acids (MUFAs), chiefly oleate (18:1) and palmitoleate (c16:1) (Enoch et al., 1976; Ntambi et al., 2002). Because this reaction is catalyzed by the *scd* gene product Stearoyl-CoA desaturase, the expression level of *scd* gene and MUFA content reflect levels of active lipogenesis. We found mRNA expression of *scd* to be enhanced in Pachón cavefish liver samples compared to surface fish after feeding (Fig. 2E). Using available lipidomics data (Medley et al., 2020), we compared the abundance of MUFAs in cavefish. We found that both oleic acid and palmitoleic acid were present in higher levels in refed Pachón cavefish livers compared to surface fish samples and cavefish samples starved for 30 days (Fig. 2F). A further indicator of lipogenesis is the fatty acid desaturation index, the ratio of product (16:1n-7 and 18:1n-9) to precursor (16:0 and 18:0) fatty acids (Chong et al., 2008; Harding et al., 2015; Klawitter et al., 2014). Indeed, we found that refed Pachón cavefish have a higher desaturation index for palmitoleic acid and oleic acid than surface fish (Fig. 2G). Taken together, this data strongly suggests that Pachón cavefish have enhanced lipogenesis in the liver.

### Pparγ is upregulated in cavefish

We checked the expression of transcription factors known to regulate lipogenesis to identify whether these may be involved in the upregulation of lipogenesis gene expression observed in cavefish. We found no significant difference in expression of the genes coding for the transcription factors Srebp1, Chrebp, Lxr, and Usf (Usf1 and Usf2) between the surface and Pachón samples (Supplementary Fig. 4). However, we noticed the gene *pparγ*, encoding a transcription factor known to be a key regulator of adipogenesis and lipogenesis (Ahmadian et al., 2013; Lee et al., 2012; Schadinger et al., 2005; Sharma and Staels, 2007; Tontonoz and Spiegelman, 2008), to be significantly upregulated in Pachón cavefish samples at both the fasted and the refed state (Fig. 3A). To test whether the differences in gene expression translate to the protein level, we generated an antibody against Pparγ. To test its specificity, we cotransfected either surface fish or cavefish *pparγ* along with GFP in HEK293T cell lines and immunostained the cells. Pparγ localized only in the nuclei of cells that were positive for GFP as the transfection control, suggesting specificity of the antibody. (Supplementary Fig. 5). We next used the antibody to quantify Pparγ protein levels in *Astyanax mexicanus*. We found higher levels of Pparγ in the liver of Pachón cavefish compared to surface fish (Supplementary Fig. 6A, B). To visualize cellular distribution of Pparγ, we performed immunofluorescence staining on liver sections. We found that Pparγ was mainly expressed in the nucleus and again found visibly higher levels in Pachón cavefish hepatocytes compared to surface fish liver cells (Fig. 3B, C). Together, these results show that Pparγ is upregulated at the mRNA and protein levels in the liver of Pachón cavefish compared to surface fish.

**Figure 3.**
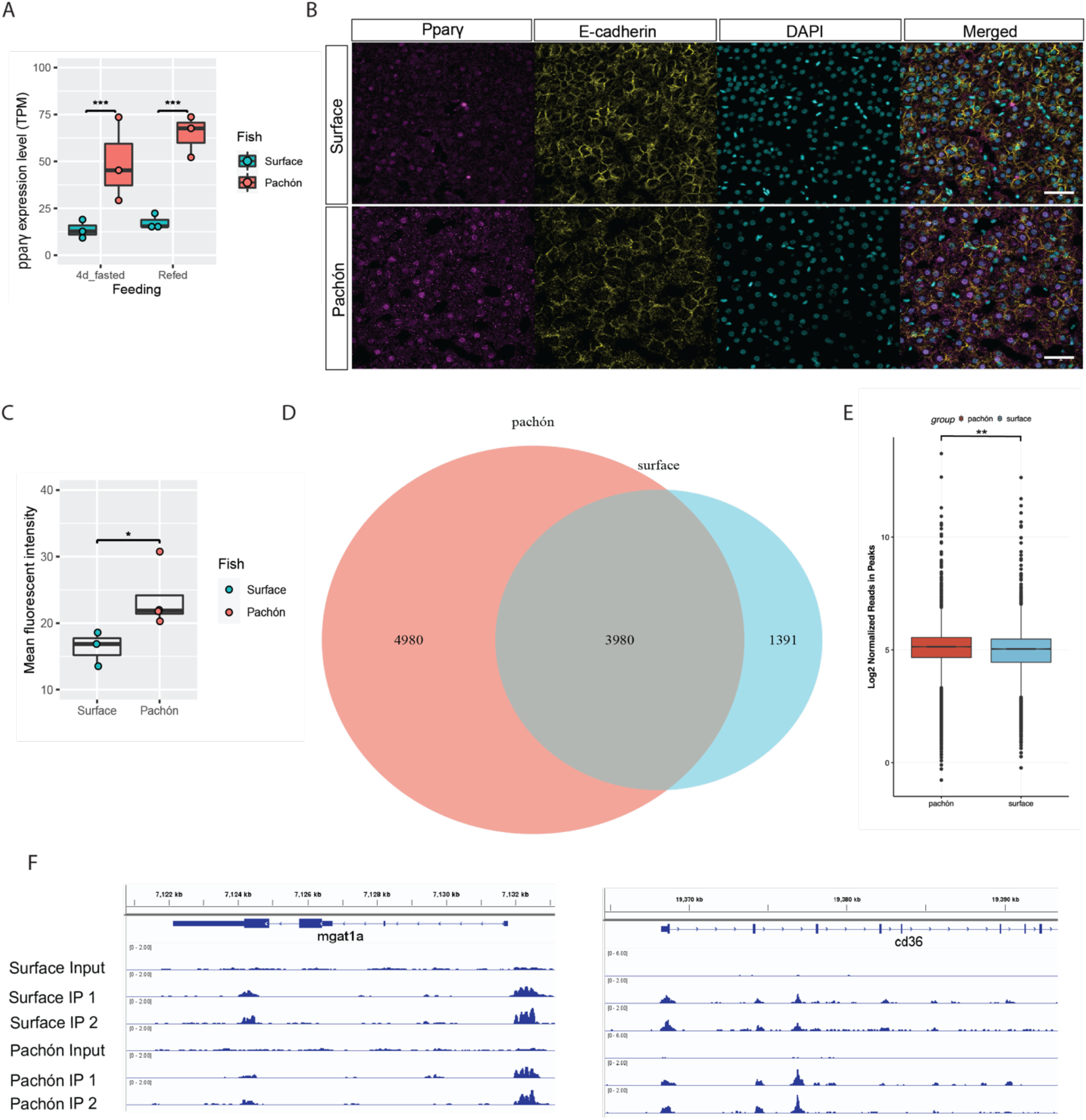
*pparγ* transcripts and Pparγ protein is upregulated in cavefish livers. A) *pparγ* mRNA expression level comparison between surface fish and Pachón cavefish under two feeding conditions: 4 day fasted and refed. TPM indicates transcript per million reads (n=3 for each group, ***p<0.001). B) Immunostaining of Pparγ (magenta), E-cadherin (yellow), and DAPI (turquoise) in liver sections of surface fish and Pachón cavefish. Scale bar =100 μm. C) Quantification of mean fluorescent intensity of Pparγ staining (n=3 for surface fish and Pachón livers respectively. 187-317 hepatocytes were randomly selected from each fish liver sample for intensity measurement (wilcox test, *p<0.05) D) Venn diagram of Pparγ ChIP-seq peaks within 3kb of predicted transcription start sites in surface and Pachón cavefish livers. E) Pparγ ChIP-seq peak height in log2 normalized read number (** p<0.01). F) Examples of Ppary ChIP-seq peaks for known lipogenesis target genes (*mgat1a, cd36*).

To characterize whether increased protein levels translate into higher binding at the DNA level, we performed ChIP-Seq for Pparγ in two livers of surface fish and Pachón cavefish, respectively. Pearson correlations between all samples showed high correlation between the biological replicates (Supplementary Fig. 7A). We used Irreproducible Discovery Rate (IDR) to keep peaks that occurred consistently in both replicates and identified 5,371 high-confidence peaks (q value<=0.01) located within 3kb of the predicted transcription start sites for the surface fish samples and 8,960 peaks for the Pachón cavefish samples (Fig. 3D, Supplementary Fig. 7B). 3,980 of those peaks were shared between two fish populations (Fig. 3D). To test if these peaks contain an enrichment for predicted Ppary binding sites, we searched all 10,251 peaks for the presence of the mouse Pparγ::Rxra motif using FIMO scan. We identified the predicted mouse Pparγ::Rxra motif in 576 (5.56%) peaks, compared to a maximum of 268 (2.59%) motifs when randomly placing the same peaks in the TSS regions of all protein coding genes (repeated 1,000 times), suggesting an enrichment of potential Ppary binding sites in our dataset (Fisher’s exact test, p < 1e-16). In addition to more genomic binding in Pachón liver samples, we found these peaks to be higher with significantly more reads than in the surface fish samples (Fig. 3E). Notably, we identified genomic binding in known PPARγ target genes involved in lipogenesis (*e.g*., mgat1a, cd36; Fig. 3F) (Feng et al., 2000; Lee et al., 2012; Zhou et al., 2008). These results are in line with our findings of higher levels of Pparγ in Pachón liver cells potentially driving expression of Pparγ target genes to a higher extent than in surface fish liver cells, providing an important data set of Pparγ genome binding sites for future studies.

### The regulation of Pparγ

PPARγ is a key regulator of lipid homeostasis and is activated by a variety of compounds, such as fatty acids, eicosanoids, and thiazolidinediones. These ligands act as PPARγ agonists that transcript the genes involved in glucose and lipid homeostasis (Georgiadi and Kersten, 2012; Kliewer et al., 1997; Krey et al., 1997; Varga et al., 2011). In addition to Pparγ, we investigated whether its activators (ligands) are also more abundant in the liver of Pachón cavefish compared to surface fish. Using published lipidomic profiling data (Medley et al., 2020), we found three natural activators (linolenic (C18:3), linoleic (C18:2), and arachidonic acids (C20:4)), to be more abundant in the livers of Pachón cavefish compared to surface fish livers (Fig. 4A). This is in line with the increased levels of the receptor, further indicating that higher lipogenesis in cavefish through Pparγ may be occurring.

**Figure 4.**
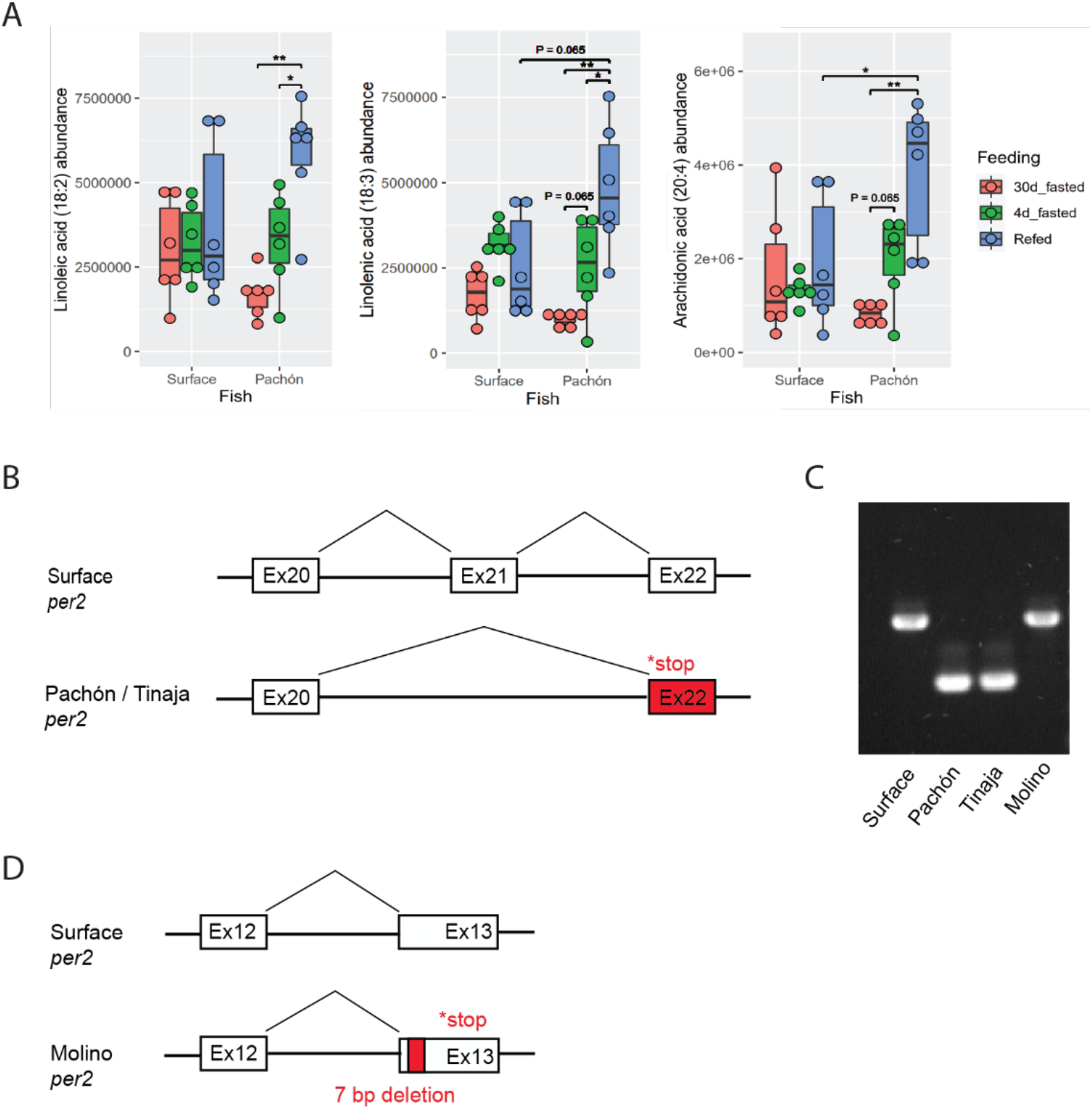
Putative regulation of Pparγ in cavefish. A) Polyunsaturated fatty acids (PUFAs), including linoneic, linolenic, and arachidonic acids, are more abundant in the liver of refed cavefish compared to surface fish liver samples (n=6 per sample, units are mTIC-normalized peak heights). Significance was calculated with Wilcoxon test *p<0.05; **p <0.01. B) Schematic depiction of the splice variant in Pachón *per2* leading to a skipping of Exon 21 and a premature stop codon in Exon 22. C) PCR with primers located in Exon 20 and Exon 22 on cDNA from liver samples, highlighting the predominance of the splice variant in Tinaja and Pachón. D) Schematic depiction of the location of the 7bp deletion in Molino *per2* Exon 13 leading to a premature stop codon in Exon 13.

Interestingly, we identified a genomic mutation in a known suppressor of Pparγ. Previously, it has been reported that Period circadian clock 2 (Per2) suppresses Pparγ-mediated transcription by direct binding to its C-terminal domain (Grimaldi et al., 2010). Analyzing the RNA-Seq liver data, we found that in Pachón liver samples, the *per2* transcript is alternatively spliced leading to a skipping of Exon 21 (Fig. 4B). The final transcript contains a premature stop codon in Exon 22, which is predicted to lead to a truncation of 160 amino acids from Per2 C-terminus in close proximity to the predicted Pparγ binding domain (Fig. 4B, Supplementary Fig. 8). We validated the splice variant using Sanger sequencing from cDNA generated from fresh liver tissue and found the alternatively spliced transcript to make up the majority, if not all, of the cDNA molecules (Fig. 4C). We also found the same splice variant to be the predominant variant in liver samples of Tinaja cavefish, but not in Molino cavefish (Fig. 4C). When we sequenced the Molino *per2* transcript, we identified a different non-sense mutation further upstream of the Pachón and Tinaja variant. Molino cavefish carry a 7 bp deletion in Exon 13 of *per2*, leading to a premature stop codon in Exon 13 and a loss of 855 amino acids from Per2, including the entire predicted Pparγ binding domain (Fig. 4D, Supplementary Fig. 8). The presence of two different non-sense mutations in *per2* in three independently derived cavefish populations indicates selection on loss of function mutations of *per2* in cavefish and a putative role in cave adaptation.

## Discussion

We sought to interrogate the cellular mechanisms contributing to high body fat accumulation in cavefish. First, we confirmed previous results showing that cavefish populations can store more fat than surface fish (Aspiras et al., 2015; Xiong et al., 2018). Our study extends previous analyses by showing that cavefish store body fat in a variety of tissues and locations in the body with certain areas more prone to body fat than others. For example, our study did not find cavefish to develop a fatty liver, which is in contrast to previous findings (Aspiras et al., 2015). This can either be due to differences in the diet of the fish used for our study, or the age of the fish used. We analyzed relatively young adult fish (~ 1 year), while previous studies have used older fish. Studying this further could provide important insights into how cavefish can deal with the accumulation of liver fat, which in humans causes nonalcoholic fatty liver disease (Rinella, 2015; Vernon et al., 2011). Furthermore, we developed a fast, reliable and cheap method of quantifying total body fat in cavefish using EchoMRI, which will open the door for high-throughput genetic analysis (i.e., QTL analysis) of fat accumulation in future studies.

Using transcriptomic analysis, we uncovered a substantial upregulation of lipogenesis enzyme genes in the liver of Pachón cavefish compared to surface fish. Moreover, the lipidomic profiling demonstrated enhanced lipogenesis level in the Pachón cavefish. In comparison to surface fish, both of these lines of evidence argue for an increased ability of cavefish to synthesize triglycerides either through de novo lipogenesis or breakdown of dietary fat. The food consumed by fish in our lab is protein rich (~60%), arguing for a high turnover through *de novo* lipogenesis, which is in line with the observed upregulation of genes involved in fatty acid synthesis (*acly*, *acaca*, and *fasn*). However, the food also contains appreciable levels of fat (~15%), which makes it likely that some of the triglyceride biosynthesis occurs through absorption and esterification of fatty acids from the dietary fat. Follow up experiments with different diets, especially high carbohydrate diets, are needed to fully disentangle this.

We also found Pparγ to be significantly upregulated in the liver of Pachón cavefish compared to surface fish. Upon ligand activation, Pparγ induces many target genes involved in lipogenesis and adipogenesis (Ahmadian et al., 2013; Lee et al., 2012; Schadinger et al., 2005; Sharma and Staels, 2007; Tontonoz and Spiegelman, 2008), making it a likely candidate transcription factor to explain the upregulation of some of the lipogenesis genes in cavefish. While Pparγ has been shown to be upregulated in obese rodent models and human patients (Lee et al., 2012; Pettinelli and Videla, 2011; Wolf Greenstein et al., 2017; Yu et al., 2003), a role of Pparγ in a species naturally adapted to food scarcity has, to our knowledge, not been reported before. Notably, we found the upregulation of *ppary* to be present already in juvenile fish (before sexual maturation) and in specific response to the feeding event, further suggesting that it has an adaptive rather than pathological role. As Pparγ plays important roles in adipogenesis, it is likely that the role of Pparγ goes beyond the increased expression of lipogenesis genes, but that Pparγ is also involved in increased adipogenesis in cavefish, potentially buffering the effect of lipotoxic lipid species (Medina-Gomez et al., 2007). Notably, we found evidence for increased genomic binding of Pparγ using ChIP-Seq analysis. However, there are some limitations to this analysis. While we have validated the specifity of the antibody in vitro, we cannot fully exclude that some of the peaks are due to unspecific binding. We did identify a highly significant enrichment for the mouse Pparγ::Rxra motif in peaks near predicted transcription start sites, however we do not know if the same motif is used in fish. Further functional analysis will be needed to fully disentangle this, however, our analysis sets an important foundation for ChIP-Seq analysis of transcription factors in non-traditional research systems.

Importantly, we identified additional signs of activation of Pparγ. Not only did we find its natural activators to be more abundant in cavefish compared to surface fish livers, but we also identified genomic mutations in one of its known repressors. Previous work has shown that PER2 represses PPARγ directly and knock-down of *Per2* leads to an increased activation of adipogenesis genes in vitro (Grimaldi et al., 2010). These findings are in line with the observed phenotypes in the cavefish. While in Molino the *per2* mutation is predicted to delete the Ppary binding domain, in Pachón and Tinaja the nonsense mutation is located 36 amino acids downstream of the predicted Ppary binding domain, potentially leaving the binding domain intact (Supplementary Fig. 8). While future experiments will be needed to explore whether these mutations affect protein structure or binding affinity for Pparγ, it is tempting to speculate that these mutations may attenuate the inhibitory effect of Per2 on Ppar*γ*-mediated transcription, which in combination with higher levels of activators would lead to higher transcriptional activity of Ppar*γ*.

Our finding of *per2* nonsense mutations in cavefish populations is interesting in terms of previous observations on this gene and circadian rhythms in general in cavefish. In studies of circadian rhythm in *A. mexicanus*, it was found that the ability to entrain a circadian rhythm is not completely lost in cavefish, but that there are differences in magnitude and timing of the rhythm (Beale et al., 2013; Mack et al., 2020). It has been speculated that this could be in part due to increased basal levels of *per2* (Beale et al., 2013; Froland Steindal et al., 2018). Our results add to these findings, potentially suggesting that Per2 is not fully functional even though its transcript is upregulated. To what extent the splice variant is present in other tissues, such as the fin or embryos, requires further investigation. However, it is clear that changes to circadian rhythm proteins are a hallmark of cavefish evolution. Interestingly, in the Somalian cavefish, *Phreatichthys andruzzi, per2* is found to have a nonsense mutation similar to the mutations we found in *A. mexicanus* (Ceinos et al., 2018) (Supplementary Fig. 8). A similar mutation in the same gene in a distantly related, convergent case of cave adaptation is an important sign of a role for this gene in adaptation. In this respect it may be worth noting that icefish have also lost *per2* (Kim et al., 2019). This is further emphasized by the fact that we found mutations in *per2* in three independently derived cavefish populations, making *per2* a major target of evolution and highlighting important connections between circadian rhythm and metabolism (Moran et al., 2014).

### Data deposition

Original data underlying this manuscript can be accessed from the Stowers Original Data Repository at https://www.stowers.org/research/publications/libpb-1619.

## Materials and Methods

### Fish husbandry

Surface, Tinaja, and Pachón morphs of Astyanax mexicanus were reared at the Stowers Institute and all animal procedures were performed with IACUC approval. The aquatic animal program meets all federal regulations and has been fully accredited by AAALAC International since 2005. Astyanax were housed in polycarbonate tanks (~2 fish per liter), with a 14:10 hour light:dark photoperiod. Each rack uses an independent recirculating aquaculture system with mechanical and biological filtration, and UV disinfection. Water quality parameters are maintained within safe limits (upper limit of total ammonia nitrogen range, 0.5 mg/L; upper limit of nitrite range, 0.5 mg/L; upper limit of nitrate range, 60 mg/L; temperature, 23 °C; pH, 7.65; specific conductance, 800 μS/cm; dissolved oxygen, >90% saturation). Standard water change rates range from 20% - 30% daily (supplemented with Instant Ocean Sea Salt [Blacksburg, VA]). A diet of *Artemia* nauplii (Brine Shrimp Direct, Ogden, Utah), Mysis shrimp (Hikari Sales USA, Inc., Hayward, CA), Gemma Micro, and Gemma Diamond 0.8 (Skretting USA, Tooele, UT) was fed to fry/juvenile/adult fish 4-12 months of age three times daily at a designated amount and directly proportional to the density of fish within the tank. The nutritional composition of Gemma Micro, according to the manufacturer, is Protein 59%; Lipids 14%; Fiber 0.2%; Ash 13%; Phospohorus 2.0%; Calcium 1.5%; Sodium 0.7%; Vitamin A 23000 IU/kg; Vitamin D3 2800 IU/kg; Vitamin C 1000 mg/kg; Vitamin E 400 mg/kg. The nutritional composition of Gemma Diamond 0.8, according to the manufacturer, is Protein 57%; Lipids 15%; Fiber 0.2%; Ash 10.5%; Phospohorus 1.6%; Calcium 2.0%; Sodium 0.5%; Vitamin A 15000 IU/kg; Vitamin D3 2400 IU/kg; Vitamin C 1000 mg/kg; Vitamin E 250 mg/kg.

Routine tankside health examinations of all fish were conducted by dedicated aquatics staff twice daily. Astyanax colonies are screened biannually for *Edwardsiella ictaluri, Mycobacterium* spp., *Myxidium streisingeri, Pseudocapillaria tomentosa, Pseudoloma neurophilia*, ectoparasites, and endoparasites using an indirect sentinel program.

### Body fat measurement

The EchoMRI™ analyzer was used to quantify fish body composition. Replicates were measured and averaged as the readout for each sample. Fat mass normalized to total body weight was indicated as body fat content. We used ~1 year old female fish of similar sizes for this and the other lipid quantification experiments.

### Total lipid content quantification

The Folch method (Flynn et al. 2009) was used to measure total lipid content. In brief, we determined dry weight by drying tissue at 60°C for 48 hours in 5 mL Eppendorf tubes (pre-weighted: W0), then measured the total weight of dried tissue and tube: W1, and calculated tissue dry weight: W1-W0. The whole tissue/organ was then homogenized with homogenizer (Benchmark Scientific, D1036) into powder. 1 mL chloroform: methanol =2:1 (v/v) was added, then samples were washed with 200 μL 0.9% NaCl. Homogenates were vortexed and centrifuged at 2,000 x g for 30 min. The lower layer (containing liquid) was transferred to pre-weighted aluminum weigh dishes (VWR, 25433-016). The liquid was dried in the hood completely (over 12h). Then, the mass of the aluminum weigh dish containing lipids was determined using a Mettler Toledo (XS105 Dual range) balance. We calculated total lipid content of the tissue using following formula: Total lipid content = total lipids (mg) / tissue dry weight (mg) * 100%.

### Hepatic triglyceride measurement

Fresh livers were collected, and mass determined. Then the hepatic triglyceride was quantified using the Triglyceride Assay Kit (ab65336) according to the manufacturer’s instructions. The triglyceride level was calculated using the following formula:

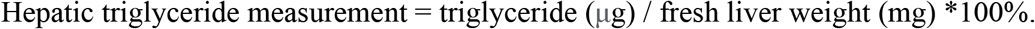

### H&E staining

The fish head and trunk were dissected and fixed in 4% paraformaldehyde for 18 h at 4 °C and embedded in paraffin while following kit instructions for dehydration, infiltration and embedding. Tissues were sectioned at 10 μm and slides were dried for 1 h in a 60 °C oven. Then, slides were stained with hematoxylin for 3 min and eosin for 1 min. Slides were washed with desalted water and air dried. Images were obtained using a VS120 virtual slide microscope (Olympus) and analyzed with ImageJ. We used ~1 year old female fish of similar sizes for this experiment.

### RNA-seq and transcriptome analysis

We used 4 months old fish for this experiment because at this stage the livers are big enough to be dissected for RNA harvest and the fish are not sexually mature, which can affect lipid metabolism heavily. Fish were housed individually for the experiment. Each fish was fed 10 mg Gemma twice per day for at least one week to allow the fish acclimate to the new environment. Once the fish were used to the new feeding regime, they were all fasted after one feeding (10 mg Gemma per fish) for 4 days. These fish were termed as the fasted group. Half of the fish (3 surface and 3 Pachón cavefish) were refed 10mg Gemma after 4 days fasting and were termed the refed group. Fish liver samples were dissected quickly, rinsed with cold PBS, and snap-frozen in liquid nitrogen. Total RNA was extracted using Trizol reagent (Ambion). Libraries were prepared according to manufacturer’s instructions using the TruSeq Stranded mRNA Prep Kit (Illumina). The resulting libraries were purified using the Agencourt AMPure XP system (Beckman Coulter) then quantified using a Bioanalyzer (Agilent Technologies) and a Qubit fluorometer (Life Technologies). Libraries were re-quantified, normalized, pooled and sequenced on an Illumina HiSeq 2500 using v4 High Output chemistry, single read 50bp, RTA v1.18.64, and bcl2fastq2 v2.20 for demultiplexing and FASTQ file generation. Both surface fish and Pachón cavefish reads were aligned to surface fish genome (Astyanax_mexicanus-2.0) via STAR aligner (v2.6.1c), under Ensembl 91 gene model. TPM gene expression values were generated using RSEM (v1.3.0). Pairwise differential expression analysis was performed using R package edgeR for different fish under different conditions. GO term enrichments were done based on upregulated and downregulated DE genes using Metascape (Zhou et al., 2019).

### RT-qPCR

The cDNA was made from 1 ug total RNA (from previous step) with high-capacity RNA-to-cDNA kit (applied biosystems, 4387406) and treated with DNase. (Promega, M6101) qPCR was conducted on a QuantStudio 6 Flex Real-Time PCR System with SYBR green detection. (Quantabio, 101414-288)). Amplification specificity for each real-time PCR reaction was confirmed by analysis of the dissociation curves. Determined *C*_t_ values were then exploited for further analysis, with the *rpl13a* gene as the reference. Each sample measurement was made in triplicate. Primer sequences for *acaca* were acaca_F 5’- CGCAGTGCCCATCTACGTG -3’ and acaca_R 5’- TGTTTGGGTCGCAGACAGC -3’. For *aclya*, the primer sequences were aclya_F 5’- GGGCACCACAGTTTTTCCAA -3’ and aclya_R 5’- CTGTCCGTGTGCCTGACTGA -3’. For *fasn*, the primer sequences were fasn_F 5’- GGGCACCACAGTTTTTCCAA -3’ and fasn_R 5’- CTGTCCGTGTGCCTGACTGA -3’. For *rpl13a*, primers were rpl13a_F 5’- GTTGGCATCAACGGATTTGG -3’ and rpl13a_R 5’ - CCAGGTCAATGAAGGGGTCA -3’.

### Fatty acid profiling

The fatty acid profiling data were extracted from (Medley et al., 2020). In brief: A group of 6 surface and 6 Pachón were starved for 30 days before dissected for liver collection (30d_fasted). A second group of 6 surface and 6 Pachón were fed regularly until 4 days before dissection (4d_fasted). A third group of 6 surface and 6 Pachón were fed regularly until 4 days before dissection. Then, on the day for dissection, they were refed 10 mg Gemma 500. Then 3 hours after they were refed, livers were collected (refed). All the livers were snap frozen and shipped to West Coast Metabolomics Center on dry ice. Fatty acids abundances were determined by charged-surface hybrid column-electrospray ionization quadrupole time-of-flight tandem mass spectrometry (CSH-ESI QTOF MS/MS). Data was reported as peak height using the unique quantification ion at the specific retention index.

### Antibody generation

The protein sequence of Pparγ was used to blast against Astyanax genomes (Astyana_mexicanus-1.0.2 and Astyanax_mexicanus-2.0) to evaluate similarity of Pparγ to other proteins in the genome. We chose 227-564aa of Pparγ as antigen for its relatively high specificity. This 338aa protein fragment was then expressed in E.coli and used to immunize two rabbits for antibody production by GenScript. ELISA titer > 1:128,000 and target protein fragment binding validation by western blot and cell line overexpression.

### Western blot

For western blot, we used four-months-old juvenile fish. The feeding regime was the same as those fish for RNA-seq. Fish liver samples were dissected quickly, rinsed with cold PBS, and snap-frozen in liquid nitrogen. Western blotting was performed using standard protocols. Briefly, liver tissues were lysed in RIPA buffer and total protein concentrations were determined by MicroBCA protein assay kit (Thermo Scientific, 23235) according to the manufacturer’s instructions and infinite 200 PRO microplate reader (Tecan). For each sample, 10 ug total protein were loaded to each well to run SDS-PAGE gel, protein transfer from gel to pvdf membrane, blocking, and antibody incubation. Imaging was carried out with Odyssey CLx system (LI-COR). The band intensity was calculated with FIJI.

### HEK293T Cell line overexpression

The surface fish and Pachón cavefish *pparγ* coding regions were cloned from cDNA, then they were inserted into pDestTol2 vector under the control of the hsp70 promoter (from zebrafish) respectively. 7.5uL FuGene (E2311) and 2.5ug plasmid were transfected into HEK293T cells on glass bottom microwell plates (MetTek, P35G-1.5-14-C). 24 hours later, 41 °C heat shock for 1 hour was performed. 48 hours after transfection, cells were fixed with 4% pfa for 20 min at room temperature (RT). Cells were permeabilized with PBST (0.1% Triton X-100) for 30 minutes at RT. Blocking was performed with Universal Blocking Reagent (BioGenex, HK085-5K,) for 1 hour at RT. A series of anti-Pparγ antibody dilutions were used to incubate cells for 2 hours at RT. After PBST (0.1% Triton X-100) wash, cells were incubated with Alexa Fluor^®^ 568 goat anti-rabbit (Invitrogen, A-11011) and DAPI (Sigma-Aldrich, D9542) for 1 hour at RT. After PBS wash, cells were imaged with Axiovert 200M microscope.

### Immunofluorescence staining

Liver tissues were fixed with 4% pfa for 16 hours at 4 °C. Then liver sections (10 μm) were done through cryostat sectioning. Slides were treated with PBS to get rid of OTC and permeabilized with 0.1% PBST (Triton X-100) for 45 min. Blocking was performed at room temperature for 1 hour. Samples were treated with TrueBlack^®^ Lipofuscin Autofluorescence Quencher (Biotium, 23007) before addition of primary antibody. Then, Primary antibody incubation was carried out at 4 °C overnight. Secondary antibody and DAPI (Sigma-Aldrich, D9542) incubation were done at room temperature for 3 hours. The antibodies in this study include anti-PPARγ (see antibody generation), anti-E-Cadherin (BD, 610182 Transduction Laboratories), goat anti-rabbit (Invitrogen, A32733), and donkey anti-mouse (Invitrogen, A31570). Images were taken with Leica TCS SP8 X microscope and analyzed with scikit-image.

### ChIP-seq

Livers from eight juvenile fish (four-month-old) were pooled together and snap frozen. Then the frozen tissues were grinded into powder, followed by 1% pfa (diluted from 16% pfa, Thermo Fisher, PI28906) fixation for 10 min at room temperature. The cross link was quenched with 0.125 M glycine. Chromatin shearing was performed by using a Bioruptor sonication system with following parameters: 30 s on and 30 s off per cycle, 10 cycles in total. DNA fragments were collected and purified with MAGnify™ Chromatin Immunoprecipitation System (ThermoFisher, 492024) according the kit instruction. Purified DNA (~10 ng) for each sample was taken as input to construct the library. Libraries were prepared using the KAPA HTP Library Prep Kit (Roche, KK8234) with 15 cycles of PCR and using 1:125 dilution of NEXTflex DNA barcodes (Perkin Elmer,NOVA-514104). The resulting libraries were purified using the Agencourt AMPure XP system (Beckman Coulter) then quantified using a Bioanalyzer (Agilent Technologies) and a Qubit fluorometer (Life Technologies). Post amplification size selection was performed on all libraries using a PippinHT (Sage Science). High throughput sequencing was performed on the Illumina NextSeq platform. Both surface fish and Pachón cavefish reads were aligned to surface fish genome (Astyanax_mexicanus-2.0). Genome browser track files in bigWig format were generated using R (version 4.0.0) packages GenomicRanges (version 1.40) (Lawrence et al., 2013) and rtracklayer (version 1.48) (Lawrence et al., 2009). Signals were normalized to fragments/reads per million (RPM). Peaks were called using MACS2 (version 2.1.2) (Zhang et al., 2008) for individual and merged replicates, respectively (q-value cutoff of 0.01). Next, IDR (https://github.com/nboley/idr, version 2.0.4.2) was used to keep those peaks that occurred consistently in both replicates. We further filtered peaks using fold enrichment and q-value cutoffs at summit position (fold enrichment ≥ 5 and q-value ≤ 1E-20). We took the summit position of the filtered peaks and used R package ChIPseeker (version 1.24.0) (Yu et al., 2015) to annotate the peaks to genomic features, including promoters (±3 Kb from transcription start site, TSS), exons, introns, downstream (within 3 Kb downstream of transcription end site), and distal intergenic regions. Astyanax genome annotation was obtained from Ensembl 98 (Yates et al., 2020). We combined and merged filtered peaks for Pachón and surface using bedtools (version 2.29) (Quinlan and Hall, 2010). For each merged peak, the new summit was assigned as the median of all overlapping peaks. Then the merged peaks were resized to 401 bp by extending 200 bp upstream and downstream of the new summits. The resized peaks were treated as the meta-peak list. We used FIMO (version 5.3.0) (Grant et al., 2011) to scan the occurrences (p-value cutoff of 1E-5) of mouse Pparg motifs (MA0065.2 in JASPAR 2020 database (Fornes et al., 2020) in 10351 meta-peaks falling into promoter regions (defined as ±3 Kb from transcription start site). To test motif enrichment, we randomly placed these meta-peaks in the promoter regions of all protein coding genes and performed FIMO scan using the same parameters. This shuffle process was repeated 1000 times.

### Per2 genotyping

Primers used to capture alternative splicing in Pachón and Tinaja (5’-3’):

Forward: CATCACTGTGACGCTCTCTCATCATCCAG

Reverse: CTCAACCAGGGATGAACCTCAGCC

PCR conditions: Denaturing 95 °C 30 sec - Annealing 57 °C 30 sec - Extension 72 °C 45 sec 35 cycles Primers used for Molino genomic DNA confirmation of 7 bp duplication (5’-3’):

Forward: CTAGGCAGTAATGATCACCTGATGAG

Reverse: GACTTGCCTGGAGCCTTTCTGGTC

PCR conditions: Denaturing 95 °C 30 sec - Annealing 56 °C 30 sec - Extension 72 °C 60 sec 35 cycles

## Acknowledgements

We thank Diana Baumann, Zachary Zakibe, Alba Aparicio Fernandez, Andrew Ingalls, David Jewell, Franchesca Hutton-Lau, Molly Miller and Elizabeth Fritz for cavefish husbandry and support. We appreciate the support from Dai Tsuchiya, Seth Malloy, and Karen J Smith of the Stowers histology support facility, the help from Rhonda Egidy, Amanda Lawlor, Michael Peterson, and Anoja Perera of the Stowers molecular biology support facility, the assistance from Alexis Murray, Olga Kenzior, and Chongbei Zhao of the Stowers tissue culture support facility, the training from Zulin Yu, Cindy Maddera, Sean McKinney, and Richard Alexander of the Stowers microscopy support facility. We thank the members of the Rohner lab for comments on the manuscript. This work was done to fulfill, in part, requirements for SX’s PhD thesis research as a student registered with the Open University, U.K.

---

**Supplementary Figure 1.**
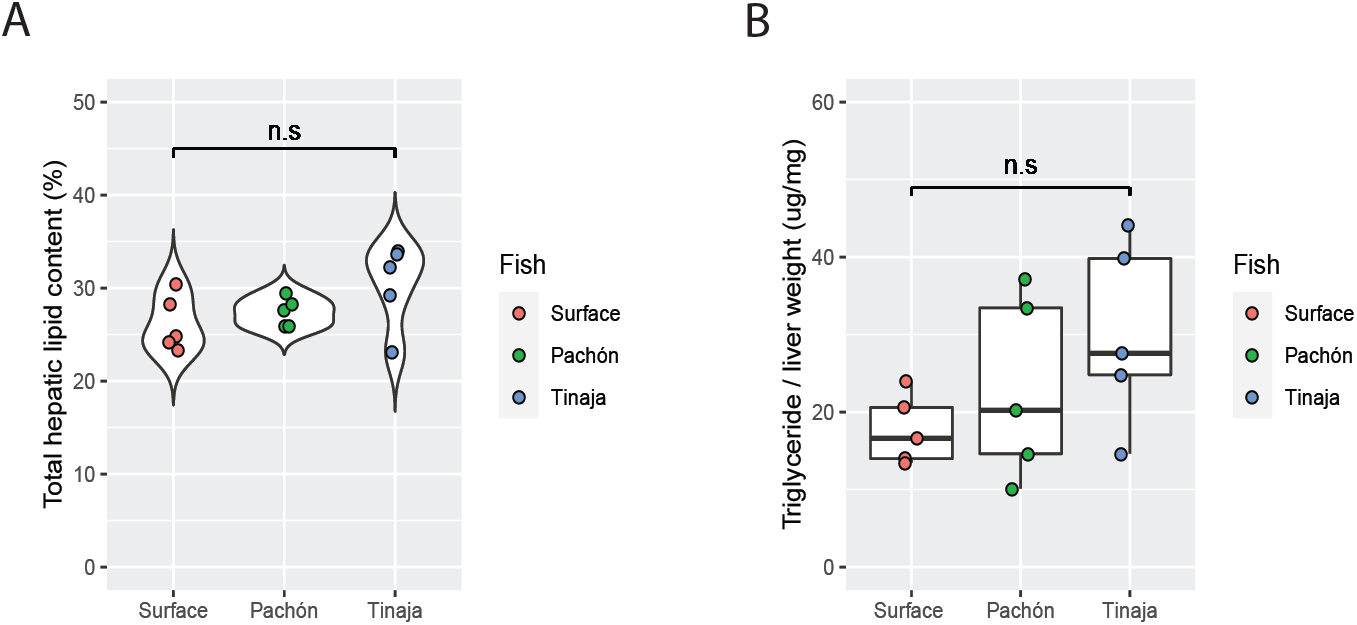
Liver fat comparison between surface fish and cavefish. A) Total lipid content (total lipid normalized to dried tissue weight) in the liver of surface fish and cavefish populations. B) Hepatic triglyceride (hepatic triglyceride normalized to fresh liver weight) comparison between surface fish and cavefish populations (n=5 per population). Wilcox test was used to determine p value and n.s indicates not significant (p>0.05).

**Supplementary Figure 2.**
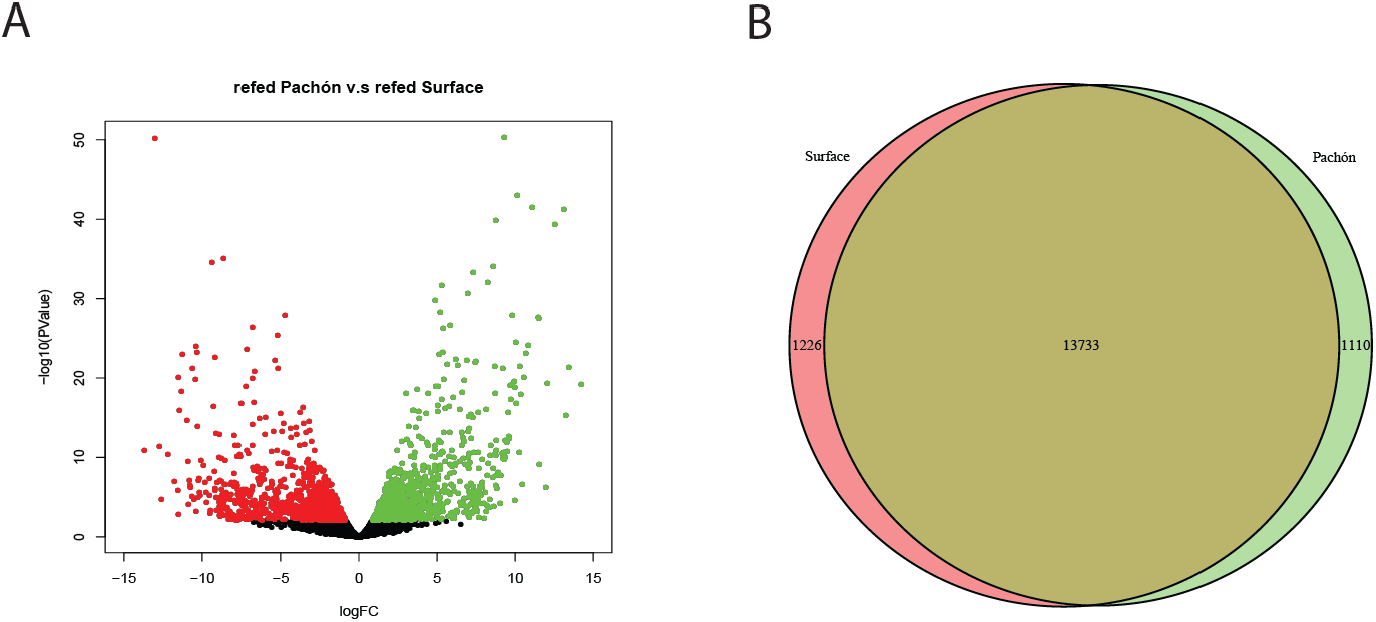
Transcriptome comparison between refed surface and refed Pachón cavefish liver samples. A) Volcano plot shows the transcriptome profile comparison between refed surface and refed Pachón cavefish. Each dot indicates one transcript. The red and green dots show the down-regulated and up-regulated transcripts in refed Pachón compared to surface fish respectively. Black dots represent transcripts with similar expression level between the two fish populations. B) Venn diagram of the transcript number comparison between refed surface and refed Pachón cavefish. From left to right: down-regulated, unchanged, and up-regulated transcripts in refed Pachón compared to surface fish respectively.

**Supplementary Figure 3.**
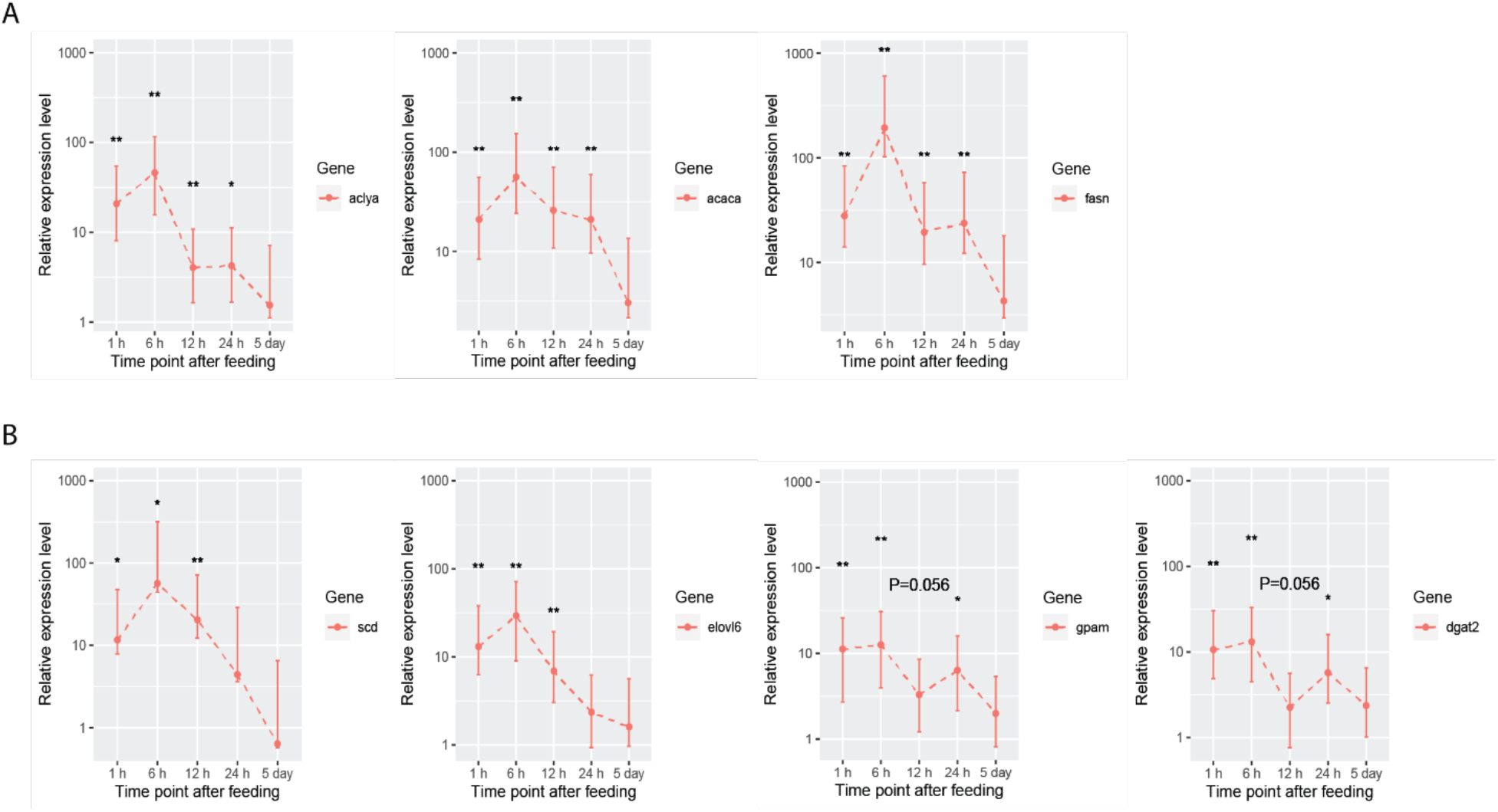
Lipogenesis gene expression dynamics after feeding. A) Relative expression level (Pachón compared to surface) change of fatty acid biosynthesis genes (*aclya*, *acaca*, and *fasn*) at different time points (1 hour, 6 hours, 12 hours, 24 hours, and 5 days) after feeding in the liver of surface fish and Pachón cavefish. B) Expression change of triglyceride biosynthesis genes (*scd1, elovl6, gpam*, and *dgat2*) at different time points (1 hour, 6 hours, 12 hours, 24 hours, and 5 days) after feeding in the liver of surface fish and Pachón cavefish. Wilcox test was used to determine p-value, (* p<0.05; ** p<0.01).

**Supplementary Figure 4.**
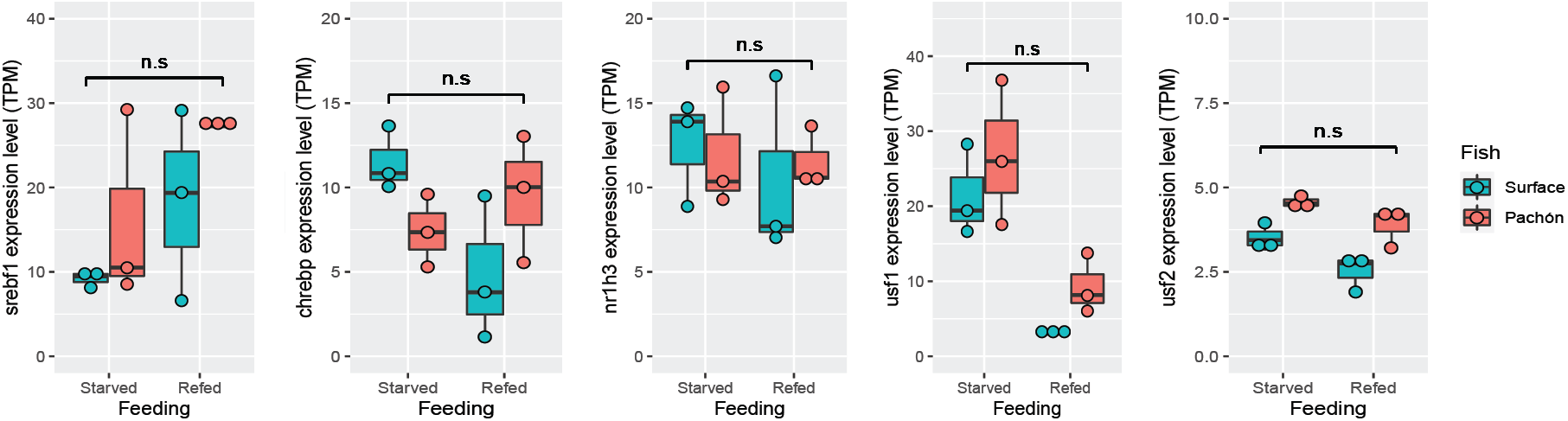
Lipogenesis transcription factor expression. RNA expression level of genes coding for known lipogenesis transcription factors in surface and Pachón cavefish livers after starvation followed by refeeding. These transcription factors include Srebf1(Srebp1), Chrebp, Nr1h3, Usf1, and Usf2. TPM means transcripts per million reads. Cyan color indicates surface fish and red color represents Pachón cavefish. Fasted and refed indicate the feeding state of the fish sample (n=3 per sample).

**Supplementary Figure 5.**
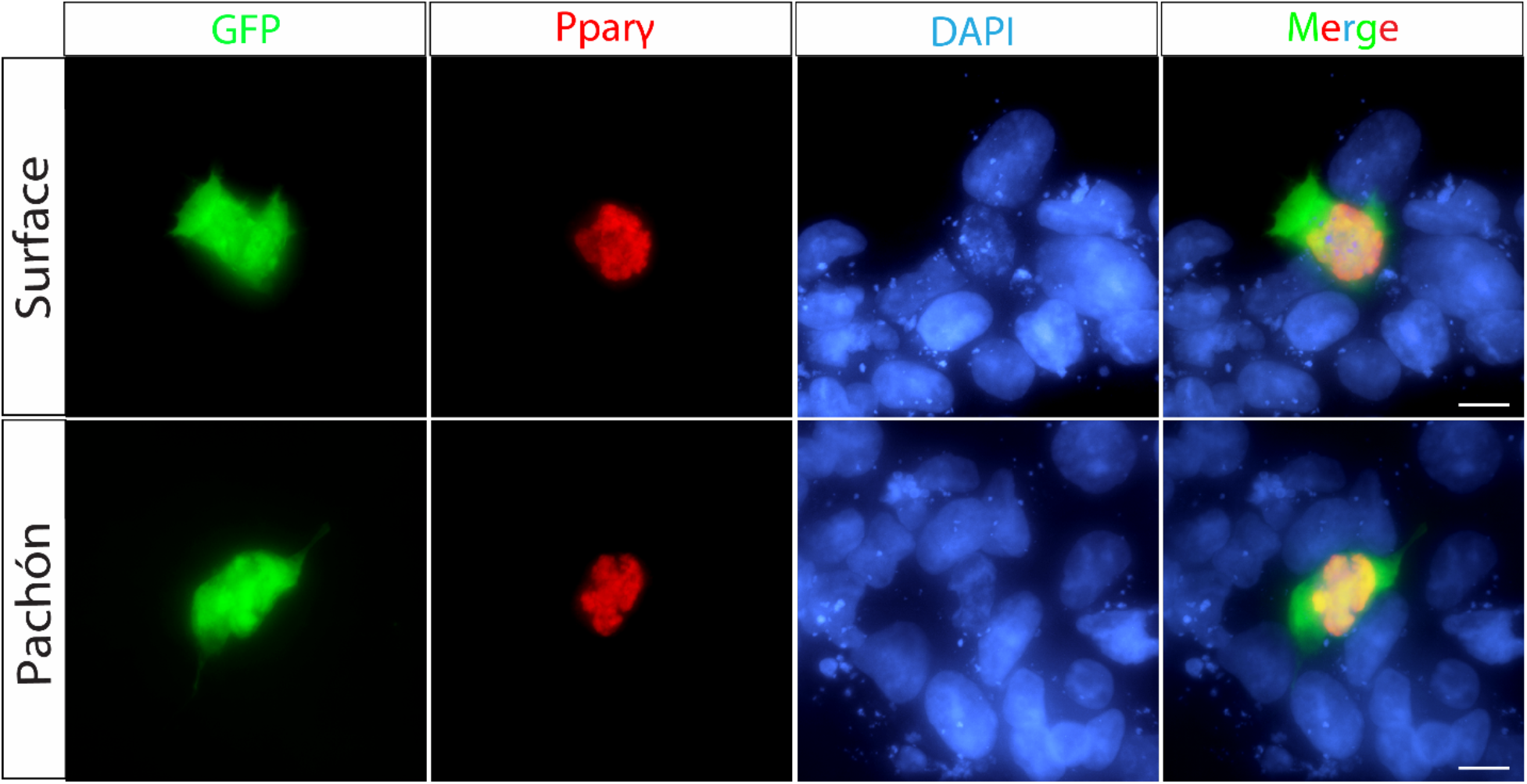
Anti-Pparγ antibody detects surface and Pachón Ppar*γ*. Transient cotransfection of expression plasmids encoding for either surface fish Pparγ (top panel) or cavefish Pparγ (bottom panel) with GFP in HEK293T cell lines. GFP (green channel) is expressed in the cytoplasm. Pparγ in surface fish or in Pachón cavefish (red channel) localizes in the nucleus. DNA was stained with DAPI (blue channel). The merge channel shows the superposition of each channel. Scale bar =10 μm.

**Supplementary Figure 6.**
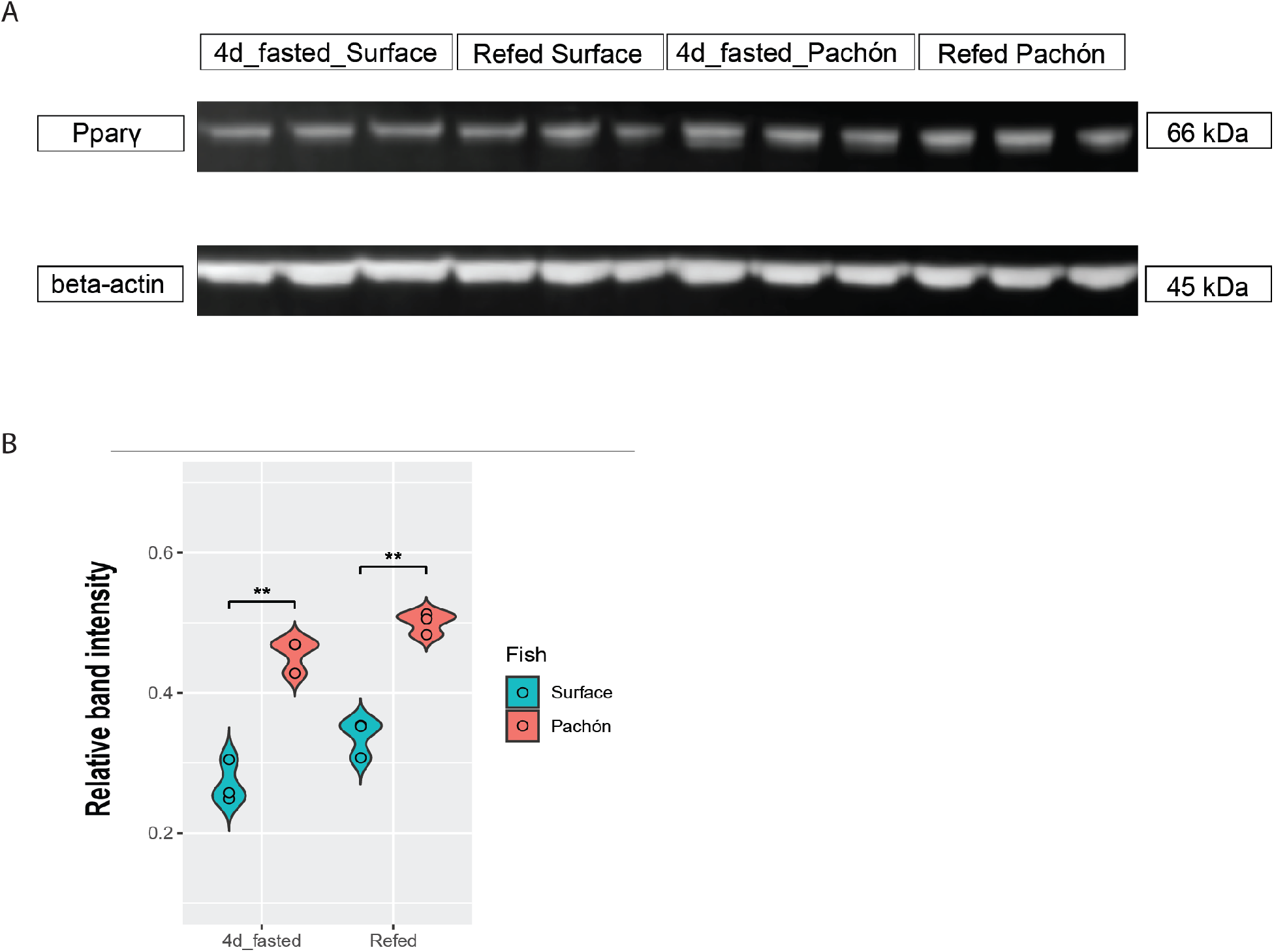
Pparγ protein expression level quantification by western blot. A) Western blots of Pparγ in the liver of surface fish and Pachón cavefish with beta-actin as loading control (n=3 for each group). B) Quantification of Western blot band intensity after normalizing to beta-actin control using ImageJ (n=3, wilcox test, **p<0.01).

**Supplementary Figure 7.**
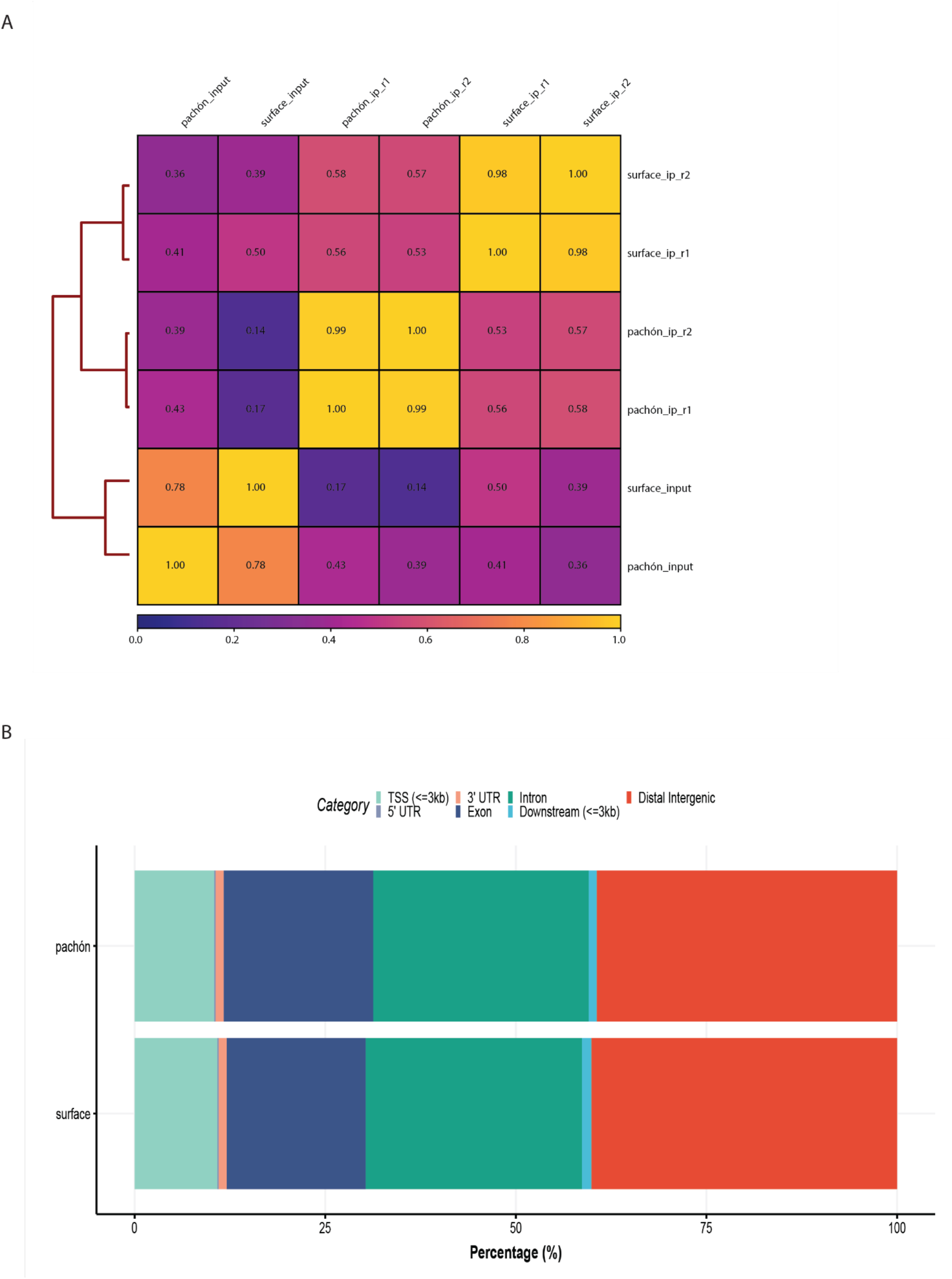
Pparγ ChIP-Seq sample correlation and genome distribution plot. A) Pearson Correlation Coefficient Sample Heatmap highlighting strong correlation between biological replicates. The correlation plot is based on the log2 read counts of bins across the genome that are generated using the R package Genomic Alignments with fixed bin size (n=10000). B) Predicted genome distribution of the peaks identified in Pparγ ChIP-Seq using the Ensembl annotation of the *Astyanax mexicanus* surface fish genome in Pachón (top) and surface fish (bottom) liver samples. For further analysis we focused on peaks near predicted TSS (<=3kb).

**Supplementary Figure 8.**
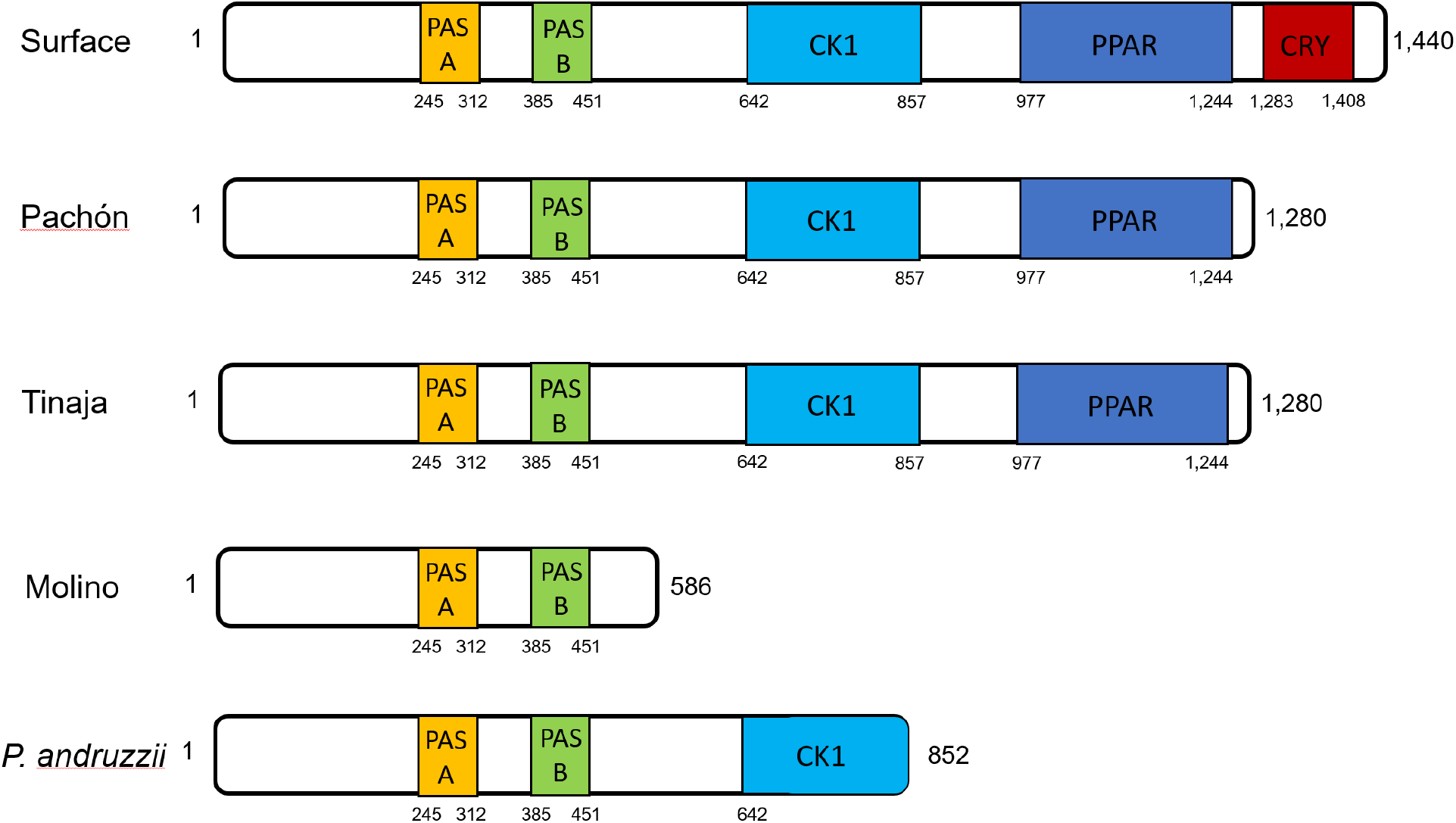
Cavefish Per2 proteins lack predicted key domains from the ancestral surface fish version. Schematic of Per2 protein with locations of Per2 mutations in *Astyanax mexicanus* cavefish populations and *Phreatichthys andruzzii* (Somalian cavefish). PPAR: homology to predicted Ppary binding domain. CRY: homology to Cry1 interacting region. PAS: homology to Per-Arnt-Sim domain; CK1: homology to casein kinase binding domain. Numbers indicate the amino acid number from N-terminal (left) to C-terminal (right).

